# Confounder-free Predictive Models for Microbiome-based Host Phenotype Prediction

**DOI:** 10.1101/2025.01.29.635502

**Authors:** Mahsa Monshizadeh, Yuhui Hong, Yuzhen Ye

## Abstract

As in many fields, the presence of confounding effects (or biases) presents a significant challenge in microbiome research, including using microbiome data to predict host phenotypes. If not properly addressed, confounders can lead to spurious associations, biased predictions and misleading interpretations. One notable example is the medication metformin, which is commonly prescribed to treat type 2 diabetes (T2D) and is known to influence the gut microbiome. In this study, we propose confounder-free predictive models for human phenotype prediction using microbiome data. These models utilize an end-to-end approach within an adversarial min-max optimization framework to derive features that are invariant to confounding factors, while accounting for the intrinsic correlations between confounders and prediction outcomes. We implemented two versions of confounder-free predictors using different network architectures: one based on a fully connected network (referred as FNN CF) and another incorporating prior biological knowledge (referred as MicroKPNN CF). We evaluated our models on microbiome datasets associated with T2D, where metformin acts as a confounder. Our results demonstrate that confounder-free predictors achieve higher accuracy compared to models that do not account for confounders and more effectively identify microbial markers associated with the phenotype, rather than markers influenced by metformin. Between the two confounder-free models, although the prior-knowledge-guided approach showed slightly lower prediction accuracy compared to the fully connected model, it offered greater interpretability, providing additional insights into the underlying biological mechanisms.

## Introduction

A fundamental challenge in biomedical research is accurately accounting for confounding variables. Confounding variables are associated with both inputs and outputs and they can obscure the true relationship between independent (input) and dependent (output) variables if not properly addressed, leading to misleading conclusions. In neuroimaging studies, for example, image data or image-derived features are often used to predict diagnostic outcomes [1]. However, if the average age of the disease group is significantly higher than that of the healthy control group—a common scenario—the age variable may act as a confounder. Without proper adjustment, predictive models may pick up on these age-related patterns, learning spurious associations rather than identifying true neurological biomarkers. In epidemiological studies, confounding variables can lead to biased results if not properly accounted for. For instance, many studies have attempted to prove a link between *Helicobacter pylori* infection and functional dyspepsia but the results have been conflicting, and it was suggested that some of the studies were biased by the selection of controls not properly matched for age, socioeconomic status and ethnic background [2].

Similarly, in microbiome research, various confounders (covariates) including host genetics, physiological status, medication use can confound the correlations between input variables and outputs such as diagnosis [3]. The risk of obtaining false or misleading results in human microbiome research is exacerbated by the substantial inter-individual heterogeneity in microbiota composition, which is likely influenced by lifestyle, physiological differences, and other factors [4]. If participants with a disease contain unique distributions of physiological or lifestyle host variables that differ from those of controls, such cross-sectional studies may conflate disease associations with the effects of confounding variables. A recent study [5] that analyzed fecal microbiota of about six hundreds of patients at different colorectal cancer (CRC) stages identified intestinal primary microbial covariates including transit time, intestinal inflammation and body mass index, which override variance explained by CRC diagnostic groups (healthy, adenoma and carcinoma). The study found that well-established CRC microbial markers [6], such as *Fusobacterium nucleatum* [7] did not significantly associate with CRC diagnostic groups when controlling for these covariates (rather, *F. nucleatum* was found to be associatied with inflammation); by contrast, potential targets including *Anaerococcus vaginalis* and *Dialister pneumosintes* remained robust [5].

Strategies can be taken to mitigate the impacts of confounders on studies that examine the role of microbiota in human diseases. There are various ways to exclude or control confounding variables including randomization, restriction (e.g., to select subjects of the same age or same sex to eliminate confounding by age sex or sex group) and matching of the subjects [8]. Matching is commonly used in case-control studies (for example, if age and sex are the matching variables, then a 50 year old male case is matched to a male control with same age). However, if these approaches are not applicable at the time of study design (impractical or impossible), researchers must rely on proper analysis of the data, using stratification or multivariate models to adjust for potentially confounding effects. For instance, by including confounders as covariates, regression models can adjust for their influence on the relationship between the independent and dependent variables [8].

Microbiome data has been widely used for host-phenotype prediction, leading to the development of various predictive models. Significant progress has been made in multiple areas, including microbiome feature extraction and the advancement of machine learning (ML) and artificial intelligence (AI) models for prediction tasks. These models differ in several aspects, such as the types of input data they use (e.g., species profiles, functional profiles, or both), the ML/AI algorithms employed, and the prediction targets (single-disease vs. multi-disease) [9, 10]. These models exhibit varying levels of prediction accuracy and interpretability—some function as black-box models, while others offer greater interpretability. Deep learning methods have been explored for representational learning of quantitative microbiome profiles in a lower-dimensional latent space, facilitating the development of predictive models. Examples include DeepMicro [11], Ph-CNN [12], PopPhy-CNN [13], and EPCNN [14]. Additionally, deep learning models capable of leveraging temporal microbiome data have been introduced. Examples include an RNN-based model for human host status prediction [15] and an LSTM-based deep learning model for food allergy prediction from longitudinal microbiome profiles [16].

We have previously developed MicroKPNN [17], which incorporates prior knowledge including phylogenetic relationship between microbial species, their metabolic activities, and microbial community information, to guide the construction of neural network in the model. We further developed MicroKPNN-MT [18], which is a multi-task microbiome-based predictor, for predicting missing metadata such as age and BMI along with disease status. What we have learned from MicroKPNN and MicroKPNN-MT is that incorporation of prior knowledge improves the interpretability of the models, and often leads to the improvements of prediction accuracy, and the model generalizability.

To the best of our knowledge, none of the existing ML/AI models explicitly considers the impacts of confounders on microbiome-based host phenotype prediction and tries to eliminate the impact of confounders on the output prediction. Motivated by the confounder-free deep learning framework proposed by Zhao et al. [1] for medical applications involving medical images (brain MRIs and X-ray images), we developed confounder-free predictors for microbiome-based predictions. We note that unlike the regression models that incorporate potential confounders as covariates, our models use the confounder information of the samples during the training process to mitigate the impacts of the confounders on the phenotypes using the adversarial optimization strategy. Once trained, the models don’t need the confounder information for predictions. This could be an advantage of using our models, as it is common that confounder information such as the metformin use are often missing. We applied our models to microbiome datasets associated with type 2 diabetes (T2D), where metformin acts as the confounder. Metformin is often prescribed to treat T2D and is known to influence the gut microbiome [19]. Our objective is to mitigate the impact of confounders (metformin in this case) on microbiome-based phenotype prediction models, thereby increasing model generalizability and reducing bias. Furthermore, we integrate metabolite-, community-, and genus-level information, allowing our model to highlight the importance of not only specific species but also the related metabolites, communities, and genera in predicting the phenotype of interest.

## Methods

Our confounder-free, end-to-end models are built on an adversarial min-max optimization framework. We implemented two versions of confounder-free predictors using distinct encoders: one based on a fully connected network (referred as FNN_CF, where CF stands for Confounder-Free) and another incorporating prior knowledge (referred as MicroKPNN_CF). In the following sections, we introduce the models, describe their training and performance evaluation, and present the datasets used for testing.

### Model architecture

As shown in Figure 1, our end-to-end models comprise three main components: an *encoder*, a *confounder predictor*, and a *phenotype predictor*. We use fully connected neural networks for the confounder predictor and the phenotype predictor. Two distinct neural network structures implement the encoders: a fully-connected structure and one incorporating a prior-knowledge primed component (so prior-knowledge informed). Figure 1 shows the encoder that incorporates prior knowledge.

**Figure 1:**
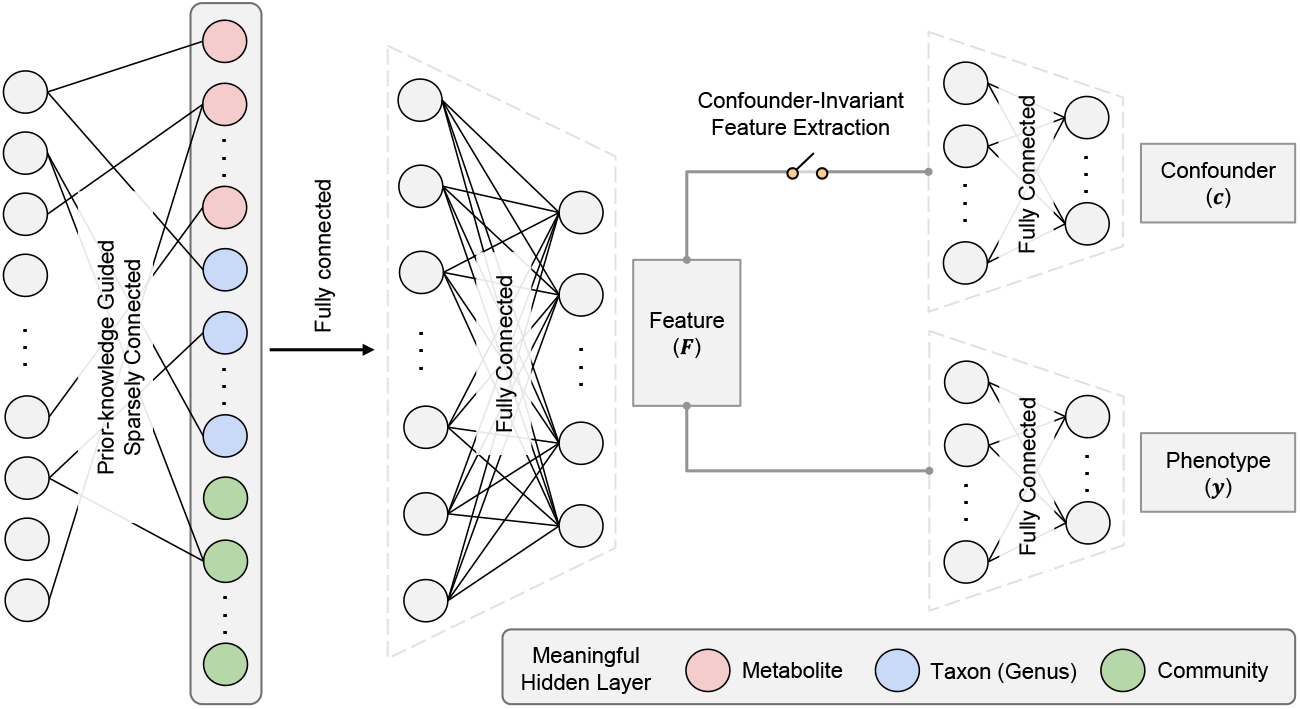
Schematic illustration of the end-to-end confounder-free models. These models contain three main components: encoder, confounder classifier, and phenotype classifier. We implemented two such models, MicroKPNN_CF, which incorporates prior-knowledge in the encoder (depicted in this diagram), and FNN_CF, which uses a fully-connected neural network for the encoder. Models are trained using the adversarial optimization approach, with the goal of maximizing the utilities of the extracted features for phenotype prediction, at the same time minimizing the utilities of the features for confounder prediction.

The encoder that leverages prior biological knowledge processes input features and explicitly models connections across multiple biological entities including metabolites that different microbial species consume and produce, taxa which reflects the taxonomic relationship between species, and microbial communities which reflect what species tend to work together. This design, as detailed in our earlier works MicroKPNN [17] and MicroKPNN-MT [18], employs a custom “MaskedLinear” layer that constrains connections between the input and hidden layers using a predefined mask. The mask encodes biologically meaningful relationships among species (in the input layer) and corresponding metabolites, taxa, and communities (in the meaningful hidden layer). Briefly, each metabolite is represented by two nodes in the hidden layer: one for production and one for consumption, and information relating producer/consumer species and corresponding nodes are derived from the NJS16 metabolic network [20]. The NCBI taxonomy is used to guide the connections between the species and their corresponding genus (for example, if two species that belong to a genus are identified in the microbiome data set, two edges will be added between the two species and the corresponding genus). Lastly, microbial communities derived from a co-occurrence network constructed from extensive microbiome datasets [21] are used to guide the connections between species and the microbial communities they belong to.

### Adversarial training

The goal of the adversarial training of confounder-free models is to derive features that are invariant to confounding factors, while accounting for the intrinsic correlations between confounders and prediction outcomes. Mathematically, the goal is to derive a feature representation **F** from input data **X** that effectively predicts the target phenotype **y** while mitigating direct influence from confounding variables **c**. To achieve this, we use an adversarial training paradigm that integrates the three components (as illustrated in Figure 1): the feature encoder, the phenotype classifier, and the confounder classifier. This setup ensures a balance between predictive performance for **y** and independence of **F** from **c**.

#### Model components

Our architecture includes the following modules:

- Encoder (*E*), a neural network *E*(**X**; *θ*_*E*_) that maps **X** to **F**, the latent representation. The encoder is optimized to maximize the discriminative capacity of **F** for *y*, while minimizing signals associated with **c**.
- Confounder classifier (*D*), the classifier *D*(**F**; *θ*_*D*_) that predicts **c** from **F**. The confounder classifier is trained using a binary cross-entropy loss with logits (*ℒ* _*c*_).

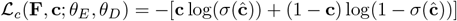

where *σ* presents the Sigmoid function, and ĉ denotes the predicted confounder, ĉ = *D*(*E*(**X**; *θ*_*E*_); *θ*_*D*_).
- Phenotype classifier (*P*), the classifier *P* (*E*(**X**; *θ*_*E*_); *θ*_*P*_) that predicts the target phenotype *y*. It is trained using a binary cross-entropy loss with logits (*ℒ* _*y*_).

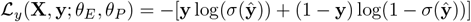

#### Training strategy

The training process alternates among three key steps to achieve confounder-resilient representation learning.

- Confounder classifier optimization: The encoder *E* and phenotype classifier *P* are held fixed, and the confounder classifier *D* is trained to minimize ℒ _*c*_. This step is performed on a subset of data where *y* = 1, ensuring that the observed variations in **F** primarily reflect differences in **c**, without interference from phenotype variations.
- Encoder debiasing: In this step, *θ*_*D*_ (confounder classifier) and *θ*_*P*_ are fixed. The encoder *E* is trained to maximize the negative correlation between **F** and the predicted confounder output using the Pearson correlation loss:

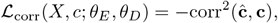

reducing residual dependence of the features on **c**.
- Phenotype classifier training: Finally, *D* is held fixed, and *E* and *P* are jointly trained to minimize ℒ _*y*_. This step is conducted over the entire dataset, ensuring that **F** retains high predictive power for **y**.

#### Adversarial optimization

The adversarial training objective integrates these steps into a min–max optimization framework:

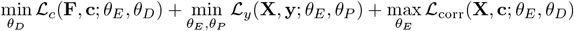

During the adversarial updates, the encoder learns to obscure confounder information that the classifier *D* could exploit.

#### y-conditioned subset and conditional independence

To enforce conditional independence **F** *⊥* **c** | **y**, the training of *D* and the encoder debiasing step use a *y*-conditioned subset *D*_*ρ*_, defined as:

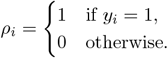

This ensures that variations in **F** during these steps are attributable solely to **c**, preserving any phenotype-mediated relationships.

### Model training, evaluation and comparison

For the confounder-free models, a learning rate of 1 *×* 10^*−*4^, batch size of 64 and latent dimension of 64 were selected based on empirical experiments considering performance and computational efficiency. For both the *D* and *P* classifiers, we used one single fully connected hidden layer, while for the *E* classifier, we utilized a knowledge-primed layer as the first layer, followed by one fully connected layer. A maximum of 100 epochs was applied, with an early stopping criterion based on the validation set performance. Specifically, early stopping was triggered if the validation balanced accuracy did not improve for 20 consecutive epochs. The optimizer Adam was used to iteratively refine *E, P*, and *D* in each epoch as described above. This configuration ensured robust performance while preventing overfitting.

All models were trained using a five-fold cross-validation scheme with stratified sampling to maintain class distributions, in which the evaluations on the validation data were used for selecting the hyperparameters. The trained models were then tested using leave-aside dataset (see below). To evaluate the models, we report metrics that account for class imbalance, including balanced accuracy (the average of recall, true positive rate, across all classes), F1 score (the harmonic mean of precision and recall), and area under the precision-recall curve (AUPRC). Results are averaged across five experiments to assess generalizability and robustness. This approach ensures that the learned representation **F** accurately predicts *y* while minimizing confounder influence.

To evaluate the effectiveness of the our confounder-free models, we compared their performances against three baseline models that we built based on Support Vector Machine (SVM), Random Forest, a Fully Connected Neural Network without consideration of confounders (FNN for short), respectively. In addition, we compared the performance of our new models against MicroKPNN [17], our previously developed model that uses prior-knowledge but without consideration of confounders. Hyperparameters for all models were optimized via grid searches on validation splits.

### Model interpretation

To interpret the trained models, we utilize DeepSHAP from the `shap` library [22] to identify the most influential input features contributing to the model’s predictions. Additionally, given that the first layer in MicroKPNN_CF is knowledge-primed with biological hierarchies, we apply DeepSHAP at this layer to assess the contributions of specific nodes, such as metabolites, taxa, or microbial communities, to the final predictions. This dual-level analysis enables a comprehensive understanding of feature importance at both the input and hierarchical levels.

The interpretability analysis is conducted for each of the five models obtained from the 5-fold cross-validation procedure. To ensure robustness and consistency, importance scores from each fold are averaged, yielding final feature importance estimates. These averaged scores provide a reliable representation of the features and nodes driving the model’s predictions, facilitating biologically meaningful insights into the underlying data.

### Datasets

Metformin is often prescribed to treat T2D and is known to influence the gut microbiome [19], and research has shown that metformin is a confounder for T2D studies [23]. As such, it is important to study the impact of metformin on training and evaluating the performance of microbiome-based predictive models. Here we utilized the microbiome data from the MetaCardis cohort [24], focusing on T2D samples and healthy controls. We derived the microbiome profile data and metadata of these samples from zenodo (record # 6242715) [25]. In total, the dataset includes 814 samples collected from individuals from three countries: France, Germany, and Denmark. For this study, samples from two countries (France and Germany) were used for training the models using 5-fold cross-validation, and samples from Denmark were used as the test set. This setting is useful for testing the generalizability of the predictive models across cohorts.

The test set consists of 143 samples, including 99 healthy controls and 44 T2D cases. The training set comprises 671 samples, with 495 T2D cases and 176 healthy controls. To prepare the data for the knowledge-primed neural network, we removed microbial operational taxonomic units (MOTUs) that were unidentified or lacked taxonomic names. Additionally, species with zero abundance across all samples were excluded. After preprocessing, the final dataset consisted of 371 features.

Figure 2 provides an overview of the distribution of samples in the training and test sets, including the proportion of T2D and control samples and whether individuals were on metformin.

**Figure 2:**
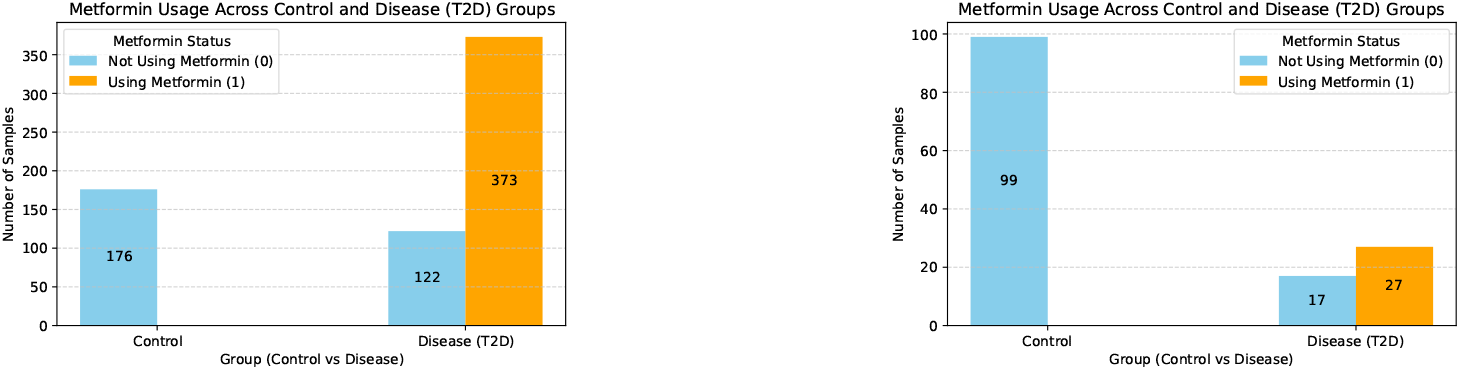
Sample distribution in the train (left) and test (right) data sets.

Data preprocessing was conducted in Python (version 3.9) using NumPy, pandas, and scikit-learn for feature transformation and metadata alignment. We applied a logarithmic transformation (log(*x* + 1)) to the microbiome abundance profile, aiming to stabilize variance, and used StandardScaler to standardize features. Log normalization has shown to be particularly useful when dealing with skewed or highly variable microbiome data, as it helps to mitigate the impact of extreme values and achieve a more balanced distribution of feature values [26].

### Availability

The proposed confounder-free (CF) framework was implemented in `PyTorch` (version 1.10) with CUDA support for efficient training on GPUs. The complete implementation, including the CF framework, preprocessing scripts, and benchmarking models, is publicly available at https://github.com/mgtools/MicrobiomeCF. The repository includes the source code, instructions for installation, and guidelines for using the framework, enabling reproducibility and adaptation to other datasets or confounding scenarios.

## Results

### Model optimization

We first assessed how well the confounder free models learned during training. Figure 3 shows the learning trajectory in different metrics for MicroKPNN_CF model.

**Figure 3:**
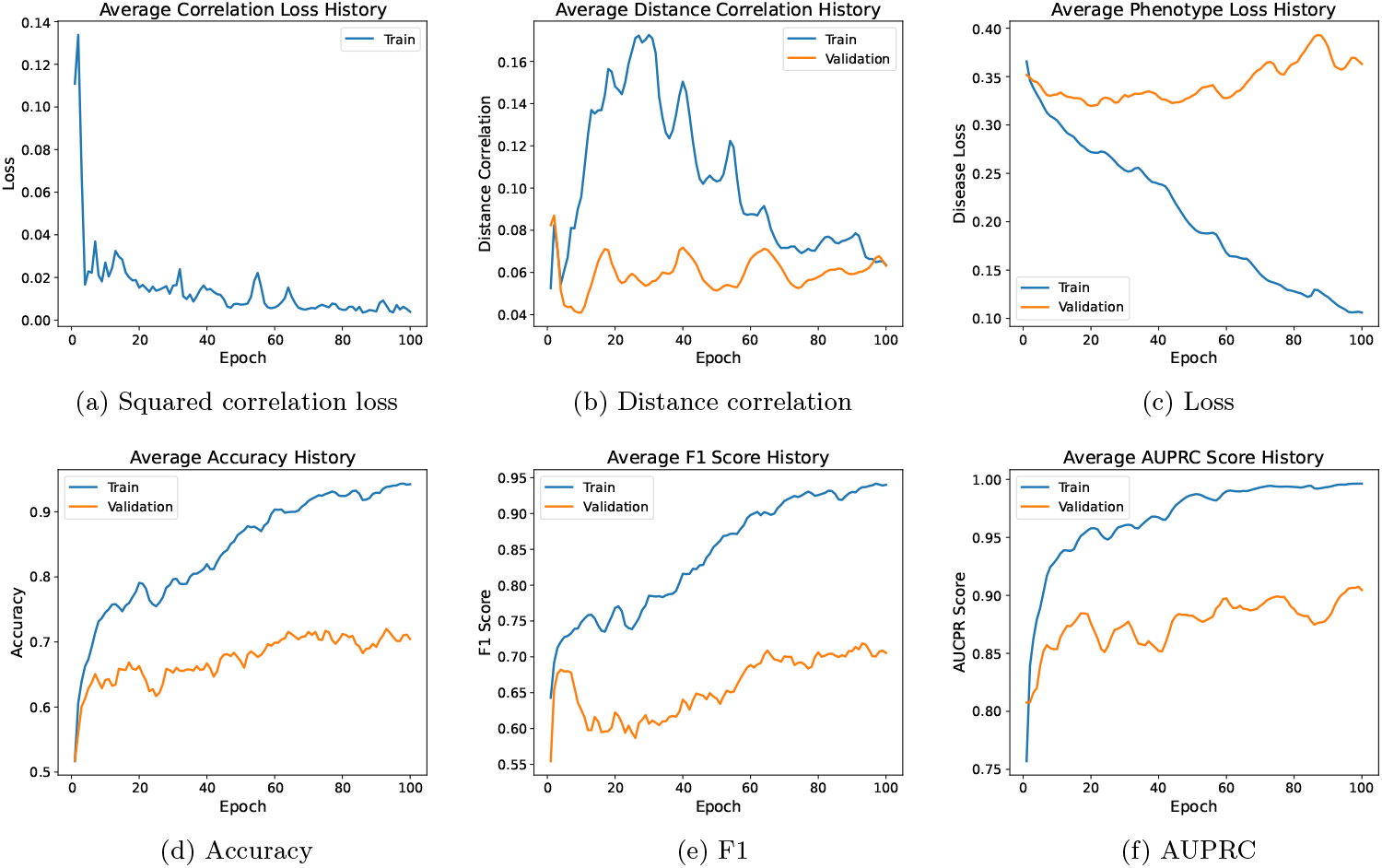
Learning trajectory of the MicroKPNN_CF model in different metrics over time. The metrices were averaged over 5-fold cross-validations.

Figure 3(a) and (c) show the trajectory of different losses that were optimized over time. Figure 3(a) shows that the squared correlation loss between the actual and predicted metformin values for individuals with the disease (-*L*_*corr*_, see Methods) decreased overtime. This indicates that the statistical mean independence between the derived high-dimensional features and the confounder metformin was reduced over time. To further ensure that the features derived from encoder are independent of metformin, we measure the statistical dependence using *distance correlation* [27], which quantifies nonlinear associations between the features and the confounder. As shown in Figure 3(b), the distance correlation initially showed a rise before gradually decreasing over subsequent epochs. This trend suggests that, in the early stages of training, the model was still learning to balance phenotype prediction and confounder independence, resulting in a temporary increase in the statistical dependence between the features and the confounder. As training progressed, the model stabilized, and the distance correlation with metformin decreased, indicating that the features became increasingly confounder-independent while preserving their relevance for phenotype prediction. Finally, Figure 3c shows the phenotype prediction loss (*L*_*y*_) over time. Together, these loss functions optimized in a min-max framework ensure that the learned features are confounder-independent and effective for phenotype prediction.

Figure 3(d)-(f) show the trajectory of accuracy over time in balanced accuracy, F1 score and AUPRC, respectively. They show that while the training accuracy kept improving, the validation accuracy reached a plateau after a certain number of epochs, indicating the importance of early stopping for better generalization of the model.

### Performance evaluation

Table 1 summaries the evaluation of the confounder-free models using the test data set in balanced accuracy, F1 score and AUPRC. We also reported the performances of other baseline methods in Table 1 for comparison. Comparing to the performances over training data set, all methods had worse performances on the test data set. RF achieved perfect predictions on training data, but its performance degraded the most on the test data set. What is encouraging is that the two confounder-free models achieved the most accurate predictions (in balanced accuracy and F1 score) comparing to the other methods. Between the two confounder-free methods, FNN_CF had slightly better accuracy. We note that SVM had a higher AUPRC on the test data set, however, AUPRC can inflate results by overfitting to the minority class, and our confounder-free models FNN_CF and MicroKPNN_CF achieve significantly higher balanced accuracy and F1 scores.

**Table 1:** Comparison of the accuracy of disease prediction by different methods (FNN_CF and MicroKPNN_CF are our confounder-free models).

| Model | Balanced accuracy (std) | F1 score (std) | AUPRC (std) |
| --- | --- | --- | --- |
| SVM (train) | 0.9463 ± 0.0033 | 0.9496 ± 0.0044 | 0.9971 ± 0.0002 |
| RF (train) | 1.0000 ± 0.0000 | 1.0000 ± 0.0000 | 1.0000 ± 0.0000 |
| FNN (train) | 1.0000 ± 0.0000 | 1.0000 ± 0.0000 | 1.0000 ± 0.0000 |
| MicroKPNN (train) | 0.9992 ± 0.0000 | 0.9992 ± 0.0001 | 1.0000 ± 0.0000 |
| MicroKPNN_CF (train) | 0.9528 ± 0.0009 | 0.9502 ± 0.0011 | 0.9973 ± 0.0000 |
| FNN_CF (train) | 0.9982 ± 0.0000 | 0.9982 ± 0.0000 | 1.0000 ± 0.0000 |
| SVM (test) | 0.6980 ± 0.0270 | 0.5944 ± 0.0230 | 0.7623 ± 0.0257 |
| RF (test) | 0.5000 ± 0.0000 | 0.4706 ± 0.0000 | 0.6048 ± 0.0430 |
| FNN (test) | 0.6755 ± 0.0015 | 0.5680 ± 0.0016 | 0.6231 ± 0.0078 |
| MicroKPNN (test) | 0.7197 ± 0.0007 | 0.6127 ± 0.0007 | 0.6804 ± 0.0036 |
| MicroKPNN_CF (test) | 0.7417 ± 0.0001 | 0.6381 ± 0.0001 | 0.6604 ± 0.0019 |
| FNN_CF (test) | 0.7530 ± 0.0001 | 0.6536 ± 0.0002 | 0.6651 ± 0.0008 |

### Confounder-resilient representation learning

As shown above, the confounder-free models (FNN_CF and MicroKPNN_CF) achieved higher accuracy of disease predictions as compared to other baseline models (SVM and RF) and neural network based models without considering the confounder (FNN and MicroKPNN).

Here we further analyzed the latent feature space *F* derived from two models FNN (without considering confounders) and MicroKPNN_CF (representing confounder-free model) to evaluate the impact of confounder mitigation strategies on representation learning. A 2D Principal Component Analysis (PCA) was employed to visualize the separation of samples based on metformin intake among T2D patients (see Figure 4) [28].

**Figure 4:**
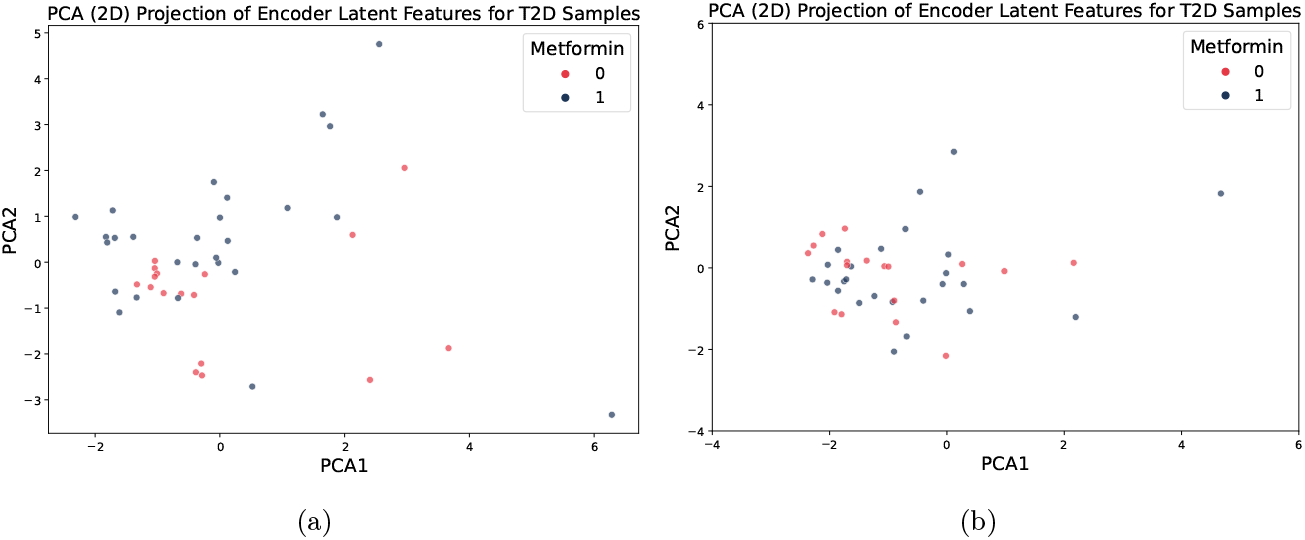
2D PCA projections of encoder latent features for T2D samples. (a) FNN-derived features *F* display distinct clusters separating patients based on metformin intake. (b) MicroKPNN_CF-derived features *F*, after confounder removal, show no clear separation between metformin and non-metformin groups, indicating that confounding influences affecting the FNN separation were mitigated.

The PCA projection for the FNN model demonstrated distinct clustering, separating metformin users from non-users. This observation was supported by a positive silhouette score of 0.063, reflecting that samples were, on average, closer to their respective clusters than to others [29]. Additionally, the within-cluster variance for the FNN model was notably low (4.98), indicating tightly grouped clusters in the latent space. The within-cluster variance metric, which quantifies the compactness of clusters, further highlights the model’s ability to create cohesive groupings [30]. These results suggest that the FNN encoder effectively captured features correlated with metformin usage, potentially influenced by the confounding factor inherent in the dataset.

In contrast, the MicroKPNN_CF model, designed with a confounder-removal mechanism, showed no discernible separation between metformin users and non-users in the PCA projection (Figure 4, right panel). The negative silhouette score (−0.087) and substantially higher within-cluster variance (20.78) highlight weaker clustering and greater overlap between the groups. This lack of distinct separation suggests that the confounding effects contributing to the clustering observed in the FNN model were successfully mitigated in the MicroKPNN_CF model. Consequently, the latent features in the confounder-free setting appeared more homogeneous, underscoring the effectiveness of the confounder-removal process implemented in MicroKPNN_CF.

### Model interpretation

We applied DeepShap to identify important input features (the species) and the important biologically-meaningful nodes in the hidden layer in MicroKPNN_CF (see Methods). We observed that comparing to baseline methods, the confounder-free models revealed species that are more likely to be associated with the phenotype (T2D) rather than the confounder. Figure 5 shows the top 20 most important species picked up by FNN and MicroKPNN_CF, respectively. The two lists shared some species (e.g., *Escherichia coli* and *Clostridium boltease* but in different ranks). Notably, FNN picked *Clostridium bartlettii* (now known as *Intestinibacter bartlettii*) as the most important species for T2D prediction, but this species is known to be impacted by the metformin use. A previous study based on a randomized trial demonstrated that metformin administration led to a decrease in the abundance of *C. bartlettii* at both 6 and 12 months of follow-up [31]. Encouragingly, MicroKPNN_CF didn’t pick it up as an important feature for predicting T2D.

**Figure 5:**
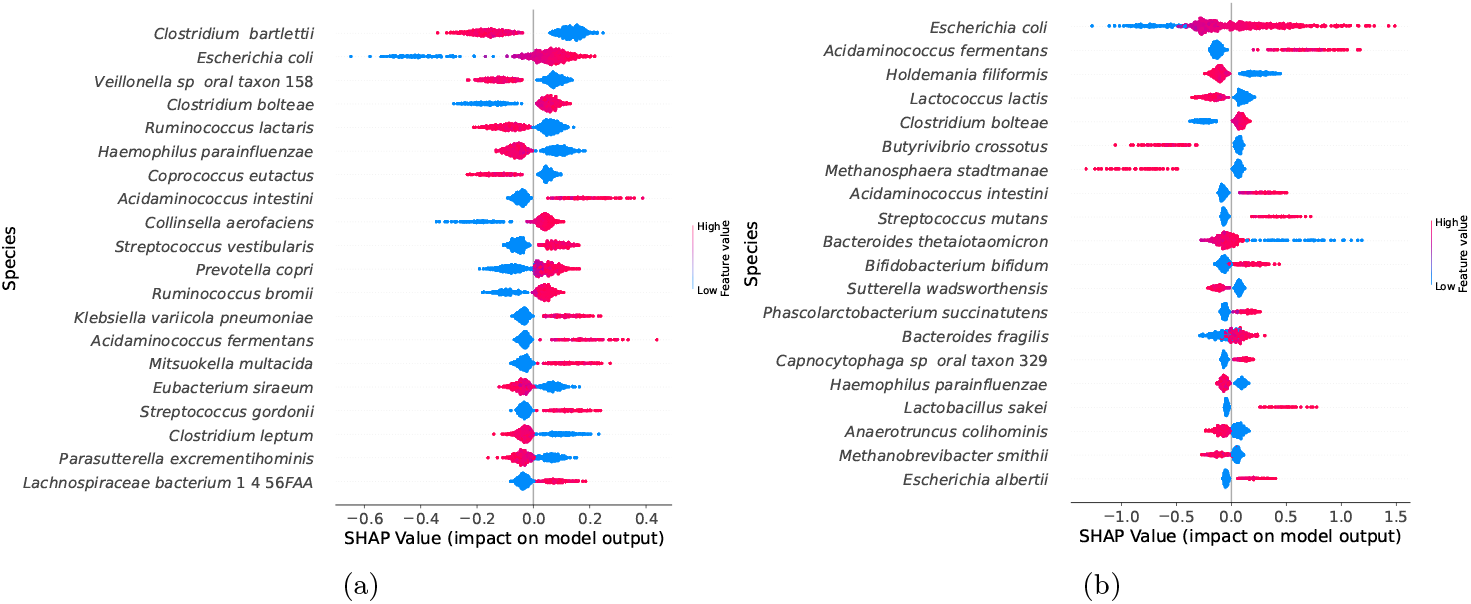
Beeswarm SHAP summary plots for comparison of important features (species) that were picked up by FNN and MicroKPNN_CF for T2D prediction. (a) FNN. (b) MicroKPNN_CF. In the plots, feature importance is ranked top-to-bottom based on average SHAP values (top = most important), and the colors indicate the feature values (red for high abundance, and blue for low abundance).

Other examples known to be associated with metformin and were picked up by FNN (which however may not be important for T2D prediction) include *Ruminococcus lactaris* [32], *Klebsiella pneumoniae* [33] and *Clostridium leptum* [34]

By contrast, important species picked up by MicroKPNN_CF were known to be associated with T2D, including *Clostridium bolteae* [35], *Butyrivibrio crossotus* [36], *Bifidobacterium bifidum* [37], *Sutterella wadsworthensis* [38], *Bacteroides fragilis* [39], and Capnocytophaga sp oral taxon 329 [40]. We note that *Escherichia coli* was picked up by both models to be important (ranked # 2 by FNN and #1 by MicroKPNN_CF). It was recently shown in a study that metformin increases gut multidrug resistance genes in T2D, potentially linked to this species [41].

Finally we showed that by incorporating prior-knowledge into the neural network, MicroKPNN_CF has the capability to reveal potential mechanism behind the important species, including the bacteria’s metabolic potential (through the metabolites), in what context (through the microbial community). Figure 6 shows the important “hidden” nodes for the predictions, including important genera (e.g., Enterocloster), metabolites (e.g., aconitic acid), and microbial community (e.g., Community 22). These observations could be used to generate hypothesis that can be examined in future research. For example, among the species that comprise community 22, *Acidaminococcus intestini* and *Megasphaera elsdenii* were previously found to be associated with T2D [42, 43]. *Acidaminococcus fermentans* is also a member of community 22, and it is a species that can consume cis-aconitate, a metabolite that was found to be important by MicroKPNN_CF.

**Figure 6:**
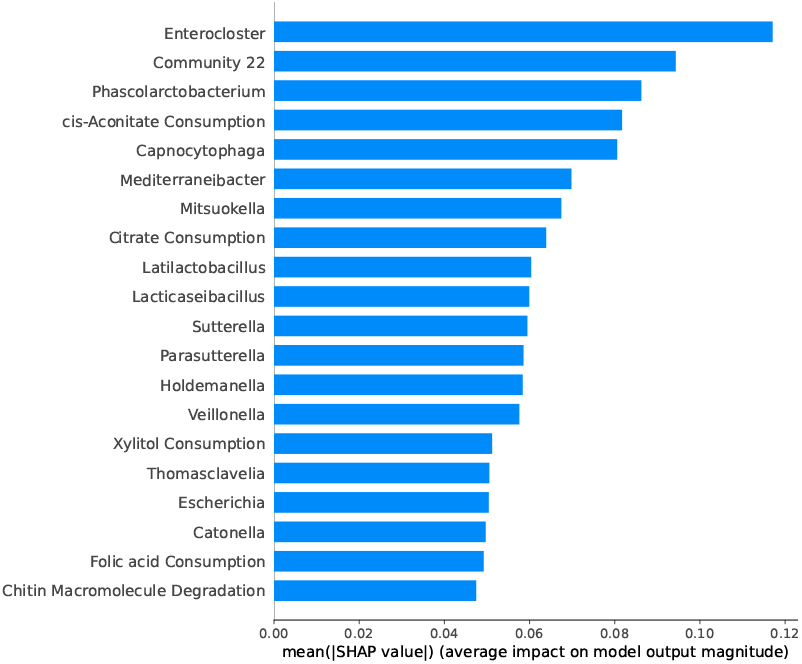
Important “hidden” nodes picked up by the MicroKPNN_CF model for T2D prediction. Here the mean SHAP values were used as the importance scores of the different nodes.

## Discussion

Our study introduces a novel confounder-free framework for microbiome-based host phenotype prediction, addressing one of the key challenges in microbiome research: the influence of confounding factors on predictive accuracy and biomarker discovery. By applying adversarial optimization techniques, we demonstrated that our models can effectively mitigate the impact of confounders, such as metformin usage in T2D studies, and improve the generalizability of microbiome-based predictive models.

The comparison between our confounder-free models (FNN_CF and MicroKPNN_CF) and traditional models highlights the importance of explicitly addressing confounders in microbiome research. Notably, FNN_CF achieved slightly higher prediction accuracy, while MicroKPNN_ CF offered better interpretability by leveraging prior biological knowledge. This trade-off underscores the value of integrating domain knowledge to uncover mechanistic insights, even at the cost of minor reductions in predictive performance.

Our results also revealed that confounder-free models are less likely to identify spurious associations caused by confounders. For instance, the FNN model ranked *C. bartlettii* as a key predictor of T2D, a species known to be influenced by metformin rather than the disease itself. In contrast, MicroKPNN_CF identified species and microbial communities with documented associations to T2D, emphasizing the biological relevance of its predictions.

Despite these advancements, the relatively lower performance on the independent test set compared to the training set highlights the challenge of heterogeneity in microbiome datasets across populations. Also, our current models only considered one confounder. Factors such as geographic, dietary, and cultural differences among cohorts contribute to variability in microbiota composition and the confounders associated with each dataset. Future studies should explore strategies to further improve model robustness, including the incorporation of additional confounders such as diet, other medications (e.g., statins or aspirin) [24], and environmental factors.

